# Toll-like receptor 7 (TLR7)-mediated antiviral response protects mice from lethal SARS-CoV-2 infection

**DOI:** 10.1101/2023.05.08.539929

**Authors:** Roshan Ghimire, Rakshya Shrestha, Radhika Amaradhi, Titus Patton, Cody Whitley, Debarati Chanda, Lin Liu, Thota Ganesh, Sunil More, Rudragouda Channappanavar

## Abstract

SARS-CoV-2-induced impaired antiviral and excessive inflammatory responses cause fatal pneumonia. However, the key pattern recognition receptors that elicit effective antiviral and lethal inflammatory responses *in-vivo* are not well defined. CoVs possess single-stranded RNA (ssRNA) genome that is abundantly produced during infection and stimulates both antiviral interferon (IFN) and inflammatory cytokine/ chemokine responses. Therefore, in this study, using wild-type control and TLR7 deficient BALB/c mice infected with a mouse-adapted SARS-COV-2 (MA-CoV-2), we evaluated the role of TLR7 signaling in MA-CoV-2-induced antiviral and inflammatory responses and disease outcome. We show that TLR7-deficient mice are more susceptible to MA-CoV-2 infection as compared to infected control mice. Further evaluation of MA-CoV-2 infected lungs showed significantly reduced mRNA levels of antiviral type I (IFNα/β) and type III (IFNλ) IFNs, IFN stimulated genes (ISGs, ISG15 and CXCL10), and several pro-inflammatory cytokines/chemokines in TLR7 deficient compared to control mice. Reduced lung IFN/ISG levels and increased morbidity/mortality in TLR7 deficient mice correlated with high lung viral titer. Detailed examination of total cells from MA-CoV-2 infected lungs showed high neutrophil count in TLR7 deficient mice compared to control mice. Additionally, blocking TLR7 activity post-MA-CoV-2 infection using a specific inhibitor also enhanced disease severity. In summary, our results conclusively establish that TLR7 signaling is protective during SARS-CoV-2 infection, and despite robust inflammatory response, TLR7-mediated IFN/ISG responses likely protect the host from lethal disease. Given similar outcomes in control and TLR7 deficient humans and mice, these results show that MA-CoV-2 infected mice serve as excellent model to study COVID-19.

## INTRODUCTION

Pathogenic human CoVs cause variable degree of disease severity that range from mild to moderate to severe respiratory illness in humans. Among the emerging pathogenic hCoVs, case fatality rate in humans caused by MERS-CoV, SARS-CoV, and SARS-CoV-2 is approximately 35%, 10%, and 1.0%, respectively (1-8). Overall, SARS and MERS epidemics each caused approximately 800 deaths, whereas SARS-CoV-2 has so far infected 700 million people with more than 6.5 million fatalities worldwide (1, 2, 7, 9). Severe COVID-19 is characterized by high fever, cough, and dyspnea with rapid progression to ALI, ARDS, and death (10-18). Patients with fatal pneumonia exhibit high lung virus loads compared to those with mild to moderate disease or those infected with a low pathogenic virus (13, 14, 16, 17, 19, 20). Virus-induced direct cytopathic effects and excessive inflammatory cytokine/chemokine responses (also known as ‘cytokine storm’) cause acute lung injury (ALI), acute respiratory distress syndrome (ARDS), and fatal pneumonia (10, 21-26).

Severe pathogenic hCoV disease, including that caused by SARS-CoV-2, is associated with excessive inflammatory cytokine activity and a simultaneously impaired type I/III IFN/ISG response (collectively referred as ‘dysregulated immunity’)(27-30). SARS-CoV-2 and other hCoVs possess multiple structural and non-structural proteins that antagonize viral RNA-induced IFN and ISG responses (reviewed in (31-34). hCoV-structural proteins (spike, nucleocapsid, membrane, and envelope) activate TLR2 and TLR4 to induce a robust inflammatory response (35-43), whereas CoVs single-stranded RNA genome and replicative intermediate double-stranded RNA elicit both antiviral and inflammatory responses (38, 44-47). CoV ssRNA primarily stimulates TLR7/8 and dsRNA mainly activates MDA5 (38, 44-47). A unique feature of the hCoV RNA is that it is abundant in GU/UU-nucleotide segments throughout its genome. These GU/UU-rich RNA are specifically recognized by TLR7/8 and induce IFN-I/III and a robust pro-inflammatory cytokine/chemokine responses (48-50) (51, 52). Conversely, dsRNA activates TLR3 and MDA5 causing robust IFN-I/III induction but a relatively weak inflammatory cytokine/chemokine expression. Several recent studies have identified the role of SARS-CoV-2 ssRNA induced TLR7 activation in IFN/ISG and robust inflammatory cytokine response (50, 53). The differential effects of TLR7/8 activation on antiviral and inflammatory response appear to be cell-type-specific. For instance, plasmacytoid dendritic cells (pDCs) produce relatively high levels of IFNs compared to monocytes-derived DCs (mDCs) or monocyte-macrophages, whereas mDCs and monocyte-macrophages are efficiently express inflammatory cytokines/chemokines (53, 54). However, the relative contribution of TLR7 induced antiviral and inflammatory response in SARS-CoV-2 pathogenesis and their impact on disease outcome are not well understood.

Since hCoV ssRNA is abundant during active infection and ssRNA-mediated TLR7 activation induces robust inflammatory cytokine/chemokine responses and IFN/ISG response, it is critical to understand the role of TLR7 in SARS-CoV-2-induced disease severity and clinical outcome. Recent *in-vitro* studies using human and murine pDCs, mDCs, or monocyte-macrophages showed that TLR7 activation induces robust IFN/ISG and inflammatory cytokine/chemokine response by SARS-CoV-2 GU-rich ssRNA stimulated or SARS-CoV-2 infected cells (49, 50, 54). Several elegant human studies showed that inborn errors in IFN response, autoantibodies against IFNs, and deficiencies in TLR3 and TLR7 signaling were associated with severe COVID-19 (53-73). Of note, X-linked recessive TLR7 defects were observed in 1% of COVID-19 patients with severe disease, and pDCs and other myeloid/myeloid-derived cells isolated from TLR7-deficient COVID19 patients showed defects in IFN/ISG response (58). These results highlight a critical role for TLR7-induced IFN/ISG response in host protection. However, given TLR7 signaling promotes both inflammatory and antiviral responses, human studies provide correlative evidence for the role of TLR7 signaling in severe COVID-19, and the *in-vivo* role of TLR7 signaling in lung antiviral and inflammatory response is unknown, the conclusive evidence demonstrating the role of TLR7 signaling in SARS-CoV-2 pathogenesis and disease outcome is lacking. Additionally, given mice are commonly used animal models for SARS-CoV-2 pathogenesis studies, it is important to define whether TLR7 deficiency in mice replicates COVID-19 phenotype observed in humans.

The primary objective of this study is to elucidate the *in-vivo* role of TLR7 signaling in SARS-CoV-2-induced inflammatory and antiviral response and assess the role of TLR7 in disease outcome. Here, we infected wild-type control and TLR7 deficient mice on highly susceptible BALB/c background with a mouse-adapted SARS-COV-2 (MA-CoV-2). WT control and TLR7 deficient mice were evaluated for morbidity and mortality, antiviral and inflammatory cytokine expression, myeloid cell accumulation, and lung pathology. We provide conclusive evidence to show that TLR7 signaling is protective during SARS-COV-2 infection. We further show that although SARS-COV-2 ssRNA-induced TLR7 activation elicits a robust inflammatory response, TLR7-mediated antiviral IFN/ISG activity likely plays a key role in controlling SARS-CoV-2 replication and host protection.

## MATERIALS AND METHODS

### Mice, virus, and infection

Specific pathogen-free male and female control BALB/c mice used in this study were purchased from Charles River Laboratories. TLR7 deficient BALB/c mice rederived on Charles River BALB/c background are described previously (44). Mice were maintained in Animal Resources animal care facility at Oklahoma State University. All the animal studies were approved by Oklahoma State University Institutional Animal Care and Use Committee (IACUC) as per American Veterinary Medical Association guidelines. Mouse-adapted SARS-CoV-2 (MA-CoV-2) strain was procured from Dr. Stanley Perlman (74). 10–16-week-old male and female mice of similar body weight were randomly allocated to two groups (WT control and TLR7 deficient) with 4-5 mice per experimental replicate. Mice were lightly anesthetized with isoflurane and were challenged with 250 PFU(LD50) of MA-CoV-2 intranasally in 50 ul of DMEM. For the survival study, mice were weighed for 10-14 days and evaluated for clinical signs (respiratory rate, fur condition, posture/movement, and ability to eat and drink). A loss of 25% of initial weight or worsening of clinical condition were considered endpoint and mortality. Mice were euthanized by isoflurane overdose based on AVMA guidelines on euthanasia.

### Virus plaque assay and gene expression study

Mice were euthanized on 2 dpi and 5dpi. Lungs were perfused with PBS via the right ventricle with 10 ml of PBS and collected. Lung tissue was homogenized using a homogenizer (Fisherbrand™ Bead mill 4) in 1 ml of PBS. One part of the lung homogenate was used for viral titer where 200 ul of 10-fold serially diluted sample was added in duplicate to each well of 12-well plate seeded with VeroE-hACE2 and incubated in 37^0^C/5%CO2 cell culture incubator with gentle rocking every 10 minutes. After 1-hour of incubation, the inoculum was removed and washed with PBS to remove any unbound virus particles and the cell monolayer was overlayed with a 1:1 mixture of 2X DMEM (2X DMEM powder 2% FBS, 1% penicillin-streptomycin, 1% L-glutamine, 1% sodium bicarbonate, 1% sodium pyruvate, 1% non-essential amino acids) and 1.2% agarose. Plates were incubated in 37^0^C/C/5%CO2 cell culture incubator for 72 hours. After 72hrs of incubation, cells were fixed in 10% paraformaldehyde for 30 minutes to inactivate the virus. Agarose was removed and stained with crystal violet (0.1%). Viral plaques were counted, and the titer was determined. Other part of the lung homogenate was collected in Trizol (TRIZOL, Invitrogen) for RNA extraction, RNA extracted was treated with RNase-free DNase (Promega Catalog: M6101) to remove genomic DNA contamination, followed by cDNA preparation using Reverse Transcriptase (Invitrogen Catalog: 28025013). RT-qPCR was performed using PCR master mix (Powerup™ SYBR™ Green Master Mix Catalog: A25741) and real-time PCR system (Applied biosystems™ 7500 Fast Real-Time PCR system). CT values were determined after normalizing to housekeeping gene GAPDH.

### Lungs cell preparation and FACS studies

To analyze phenotype of different immune cell populations in the lungs of MA-CoV-2 infected control and TLR7 deficient mice, PBS-perfused lung tissue was treated with collagenase-D and DNAse-1. For surface staining, single cell suspension isolated from lung tissue was treated with Fc block (anti-CD16/32) followed by incubation with fluorochrome-conjugated mouse-specific antibodies: PECy7 α-CD45 (clone: 30-F11); FITC α-Ly6G (clone: 1A8, BD Biosciences); PE/PerCp-Cy5.5 α-Ly6C (clone: AL-21 [BD Biosciences] or clone: HK1.4); V450 α-CD11b (clone: M1/70). Isolated cells were surface labeled for Neutrophil (CD45^+^ CD11b^+^ Ly6G^hi^) and inflammatory monocyte (CD45^+^ CD11b^+^ Ly6c^hi^). Cells were fixed with Cytofix solution (BD Bioscience Catalog no: 554655), flow data were acquired on BD FACS-ARIA-III™ and analyzed using FlowJo™ software. Surface staining was carried out using a previously described protocol (75). All fluorochrome-conjugated antibodies were used at a final concentration of 1:200 in FACS buffer, except for PE/PerCp-Cy5.5 α-Ly6C labeled antibody which was used at 1:300 concentration.

### Statistical analysis

Statistical analysis was done using GraphPad Prism Version 9.5.1 (GraphPad Software, Inc). Results obtained were analyzed using Student’s t-test or One-way ANOVA, with bar graphs and scatter plot data represented as mean +/-SEM. Statistical significance of weight loss was determined using One-way ANOVA and significance of survival curves were determined using log-rank (Montel-Cox test) or Gehan-Breslow-Wilcoxon test. A threshold value of *P<0.05, **P<0.01, or ***P<0.001 was used to assess the statistical significance.

## RESULTS

### TLR7 signaling protects mice from SARS-COV-2-induced morbidity and mortality

Loss of function TLR7 variant or X-linked TLR7 mutation/deficiency is associated with severe COVID-19 in humans (53, 76-82). To conclusively demonstrate the role of TLR7 signaling in SARS-CoV-2 pathogenesis and disease outcome, we utilized 12–16-week-old WT control and TLR7 knockout (TLR7 deficient) mice on highly susceptible BALB/c background. Control and TLR7 deficient male and female mice were infected with LD_50_ (250pfu) of mouse-adapted SARS-CoV-2 (MA-CoV-2) via the intranasal route and monitored for 14 days to assess morbidity (percent initial weight loss) and mortality. Our results show that both control and TLR7 deficient BALB/c mice lost significant weight until day 6 p.i., while the recovery was slow in TLR7 deficient mice as compared to control BALB/c mice (Figure 1A), with approximately 75% of TLR7 deficient mice ultimately succumbing to infection compared to ∼40% mortality observed in control infected mice (Figure 1B). These results conclusively demonstrate that TLR7 signaling is critical for host protection during SARS-CoV-2 infection and support human studies that show an association between TLR7 deficiency or a loss of function TLR7 variants with severe COVID-19.

**Figure 1:**
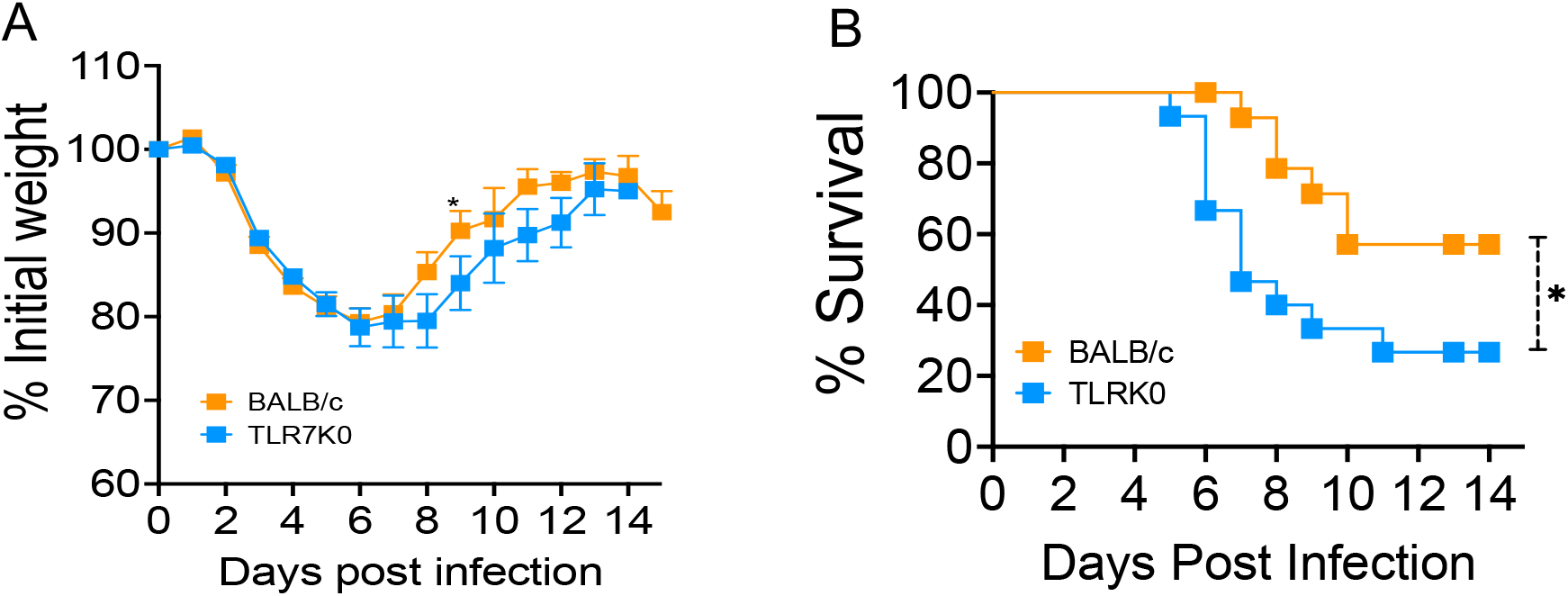
TLR7 signaling protects mice from SARS-CoV-2-induced morbidity and mortality. 12–16-week wild type BALB/c mice and TLR7KO mice were infected intranasally with 250PFU of MA-CoV-2. Percent initial weight (A) and survival (B) of mice were monitored for 14 days following infection. Data are representative of 3 independent experiments with 3-5 mice/group. Statistical significance was determined using log-rank (Montel–Cox test) or Gehan–Breslow–Wilcoxon test with \**P* < .05 and \*\**P* < .01.

### TLR7 signaling is essential for antiviral IFN/ISG induction during SARS-CoV-2 infection

We and others showed that TLR7-signaling was essential for antiviral IFN/ISG response and host protection during mouse CoV (MHV, mouse hepatitis virus) and MERS-CoV infection (28, 44, 46). Similarly, pDCc, DCs, and myeloid DCs from TLR7 loss of function variant COVID-19 patients showed reduced IFN/ISG levels compared to those from control COVID19 patients (53-55, 79, 80). However, the *in-vivo* role of TLR7 signaling in lung IFN/ISG response during SARS-CoV-2 infection is not known. Therefore, to confirm whether TLR7 deficiency leads to reduced IFN/ISG response in SARS-CoV-2 infected lungs, we examined mRNA levels of type I IFNs (IFNα/β), type III-IFN (IFNλ), and ISGs (ISG15 and CXCL-10) in naïve and MA-CoV-2 infected control and TLR7 deficient lungs. As shown in figure 2, compared to naïve mice, we observed a significant increase in IFN-I/III and ISG levels in control-infected mice at day 2 post-infection. Further comparison of IFN-I/III and ISGs levels in MA-CoV-2 infected control and TLR7 deficient mice showed a significant decrease in transcript level of IFNs (IFNα/β and IFNλ) in TLR7 deficient mice as compared to control BALB/c mice (Figure. 2A-B). These results show that similar to other hCoV infections, TLR7 is required for IFN-I/III and ISG induction in SARS-CoV-2-infected lungs.

**Figure 2:**
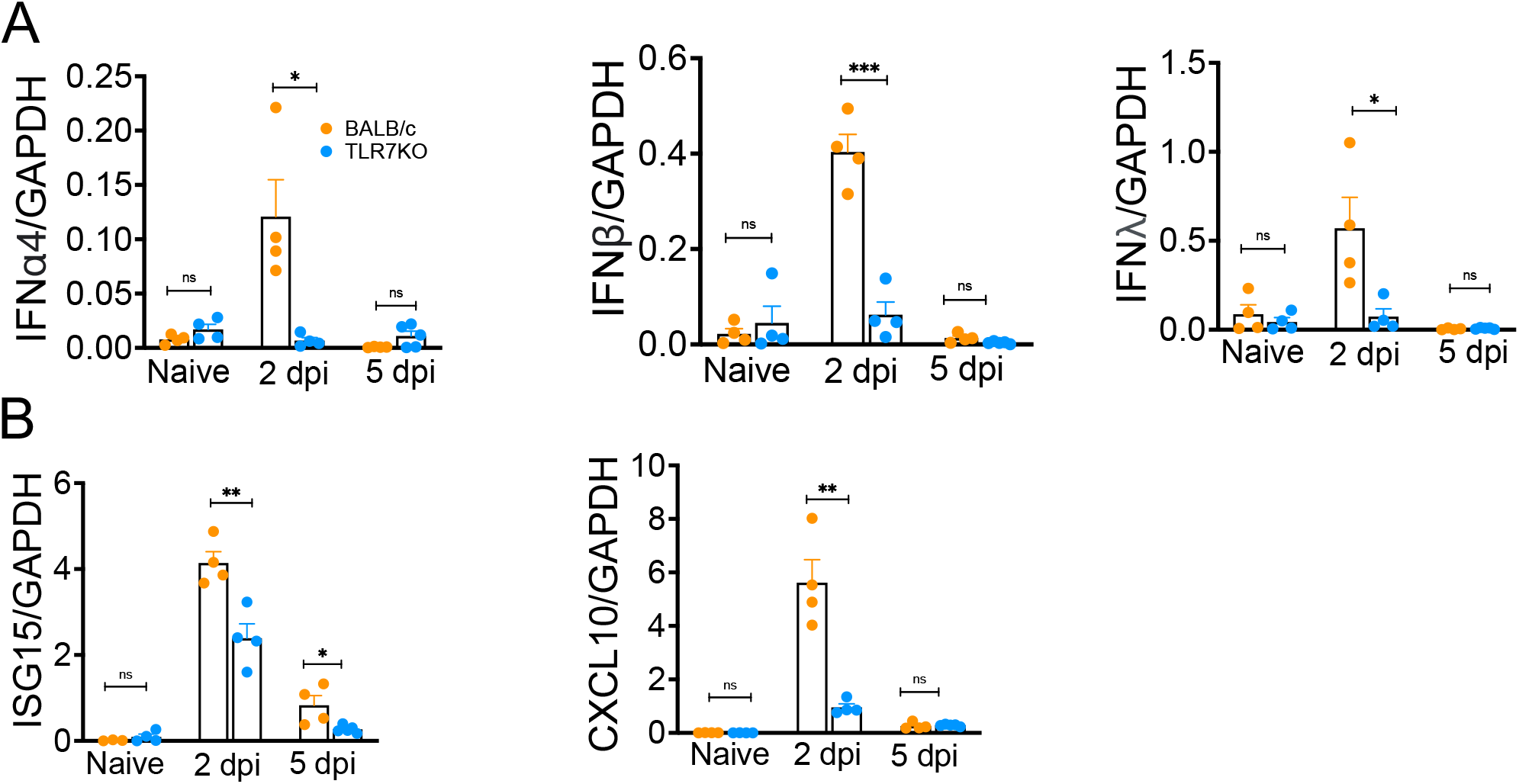
TLR7 signaling is essential for antiviral IFN/ISG response in SARS-CoV-2 infected lungs. 10-16-week wild type BALB/c mice and TLR7KO mice were intranasally infected with 250 PFU of MA-CoV-2. Lung tissue was harvested on 2 and 5dpi. Relative mRNA levels of indicated genes were determined by qPCR. (A) Graphs show relative mRNA levels of IFNs (A) and ISGs (B) in naïve and infected control and TLR7KO mice. Results are representative of two independent experiments with n=4-5 mice per group per experiment. Statistical significance was determined using Student’s t test with *P < .05 and **P < .01.

### TLR7 promotes robust inflammatory cytokine expression in SARS-CoV-2-infected lungs

Pathogenic hCoV RNA, including SARS-CoV-2 genome, is abundant in GU/UU-nucleotide sequences. These GU/UU-nucleotides specifically stimulate TLR7/8 to induce robust pro-inflammatory cytokine and chemokine production (49, 50, 83). We also showed that ssRNA mimics induce robust pro-inflammatory cytokine production compared to dsRNA and DNA mimics (75). Therefore, to examine whether TLR7 signaling is essential for robust inflammatory cytokine production in MA-CoV-2 infected lungs, we examined lung mRNA levels of several inflammatory cytokines and chemokines. Control and TLR7 deficient mice infected with 1LD50 of MA-CoV-2 were euthanized at days 2 and 5 post-infection. Quantitative PCR analysis showed significantly increased mRNA transcript levels of pro-inflammatory cytokines and chemokines (TNF, IL-6, IL-1b, CCL-2, CXCL-1) in MA-CoV-2 infected lungs compared to naïve lungs (Figure. 3). Additionally, similar to IFN/ISG expression levels, MA-CoV-2 infected TLR7 deficient lungs showed significantly reduced mRNA levels of several pro-inflammatory cytokines and chemokines compared to infected control lungs (Figure. 3). These results demonstrate that TLR7 signaling not only promotes IFN/ISG expression but also facilitates SARS-CoV-2 induced robust inflammatory cytokine and chemokine production in the lungs.

**Figure 3:**
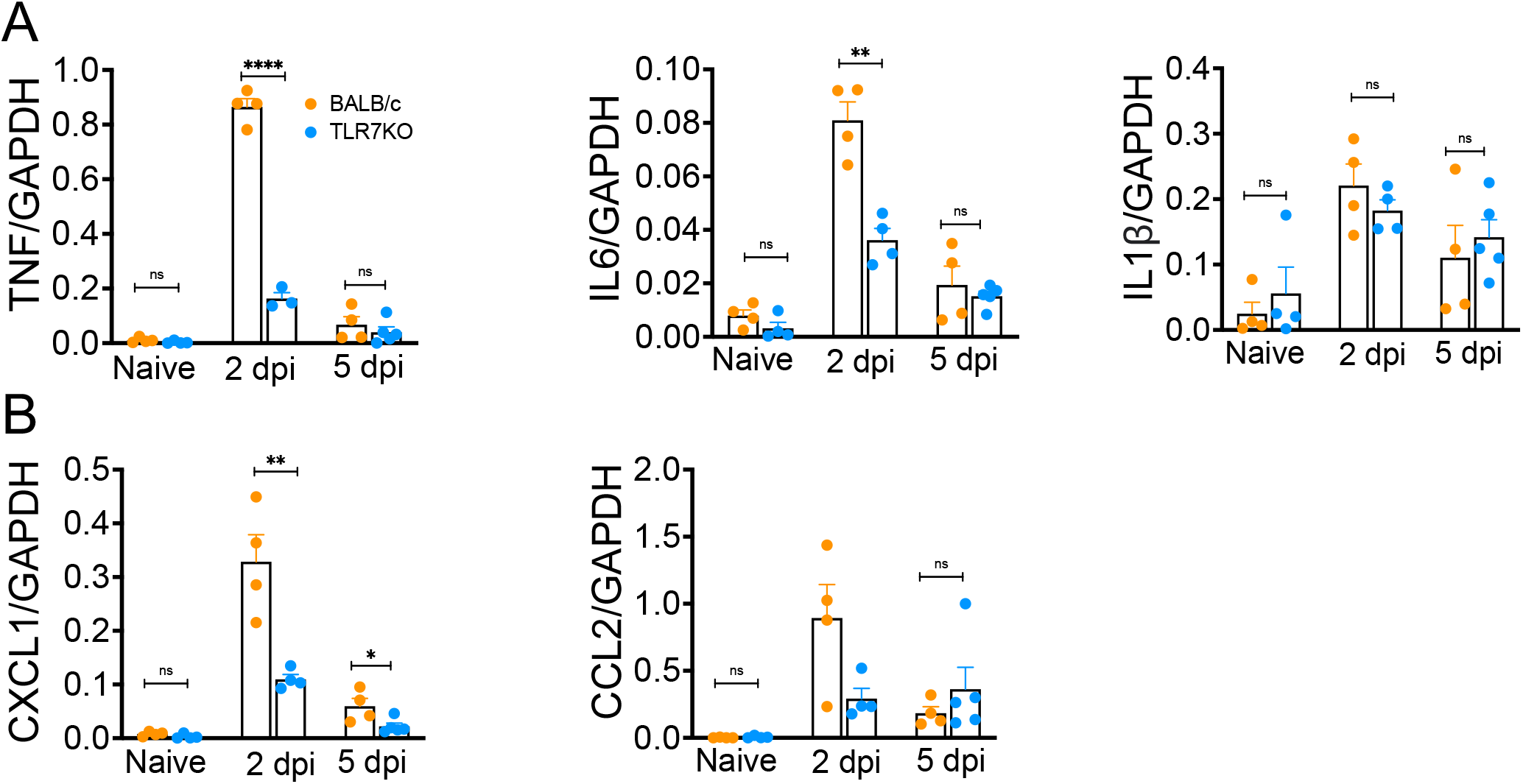
TLR7 promotes inflammatory cytokine/chemokine expression in SARS-CoV-2 infected lungs. 10-16-week control BALB/c mice and TLR7KO mice were intranasally infected with 250 PFU of MA-CoV-2. Lung tissue was harvested on 2 dpi and 5dpi. Relative mRNA levels of indicated inflammatory cytokines/chemokines were determined by qPCR. Graphs show mRNA levels of inflammatory cytokines (A) and chemokines (B). Results show one representative study of two independent experiments with n=4-5 mice per group. Statistical significance was determined using Student’s t test with *P < .05 and **P < .01.

### TLR7 activation suppresses SARS-CoV-2 replication in the lungs

Our results in figure 2 showed that TLR7 deficiency leads to reduced SARS-CoV-2-induced IFN/ISG responses. Therefore, to confirm whether reduced IFN/ISG response in TLR7 deficient mice correlates with increased MA-CoV-2 replication, we examined lung virus titers and sub-genomic RNA levels in control and MA-CoV-2 infected lungs at days 2 and 5 post-infection. As shown in figure 4A-B, we observed a significant increase in MA-CoV-2 titers and sub-genomic RNA levels in TLR7 deficient lungs compared to control mice at day 2 and 5 post-infection. These results demonstrate that TLR7-induced IFN/ISG responses are critical for suppressing SARS-CoV-2 replication in the lungs.

**Figure 4:**
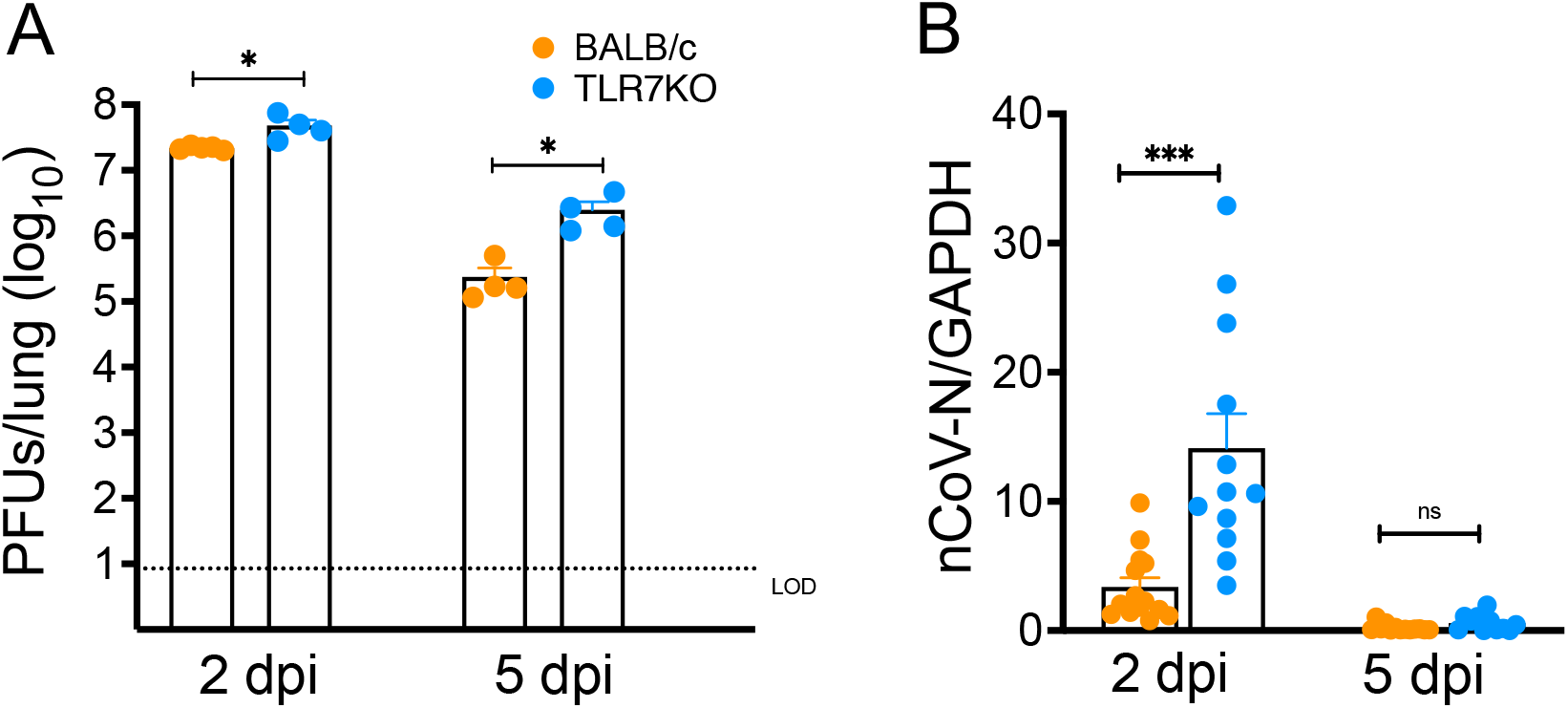
TLR deficiency results in increased virus titers in the lungs. 10–16-week control BALB/c mice and TLR7KO mice were infected intranasally with 250PFU of MA-CoV-2. Bar graph represents log_10_ change in SARS-CoV-2 titers (A) and sub-genomic mRNA level relative to GAPDH (B) in control and TLR7KO lungs. A) Data are representative or B) pooled from two to three independent experiments with 4-5 mice/group. Statistical significance was determined using Student’s t test with *P < .05 and **P < .01.

### TLR7 deficiency leads to differential myeloid cell accumulation in SARS-CoV-2-infected lungs

Excessive myeloid cell activity, specifically monocyte-macrophage and neutrophil responses, cause severe hCoV disease (26, 84-89). Robust IFN-I and inflammatory cytokine/chemokine responses drive myeloid cell influx and inflammatory (28, 44, 90). To examine whether lack of TLR7 signaling leads to differential myeloid cell response in lungs, we euthanized MA-CoV-2 infected control and TLR7 deficient mice at 2 dpi and 5 dpi. PBS-perfused collagenase/DNAse digested lungs were analyzed for myeloid cell infiltration using flow cytometry. As shown in figure 5A-D, we observed a significant decrease in the percentage of inflammatory monocyte-macrophages at 2 dpi, but the total number of these cells was not different in infected control and TLR7 deficient lungs at either day post-infection. In contrast, we found a significant increase in the percentage and total number of Neutrophils at 5 dpi (Figure. 5). These results collectively show that the loss of TLR7 signaling results in differential accumulation of myeloid cells with a specific increase in neutrophils in lungs which may contribute to increased mortality in TLR7 deficient mice compared to control mice.

**Figure 5.**
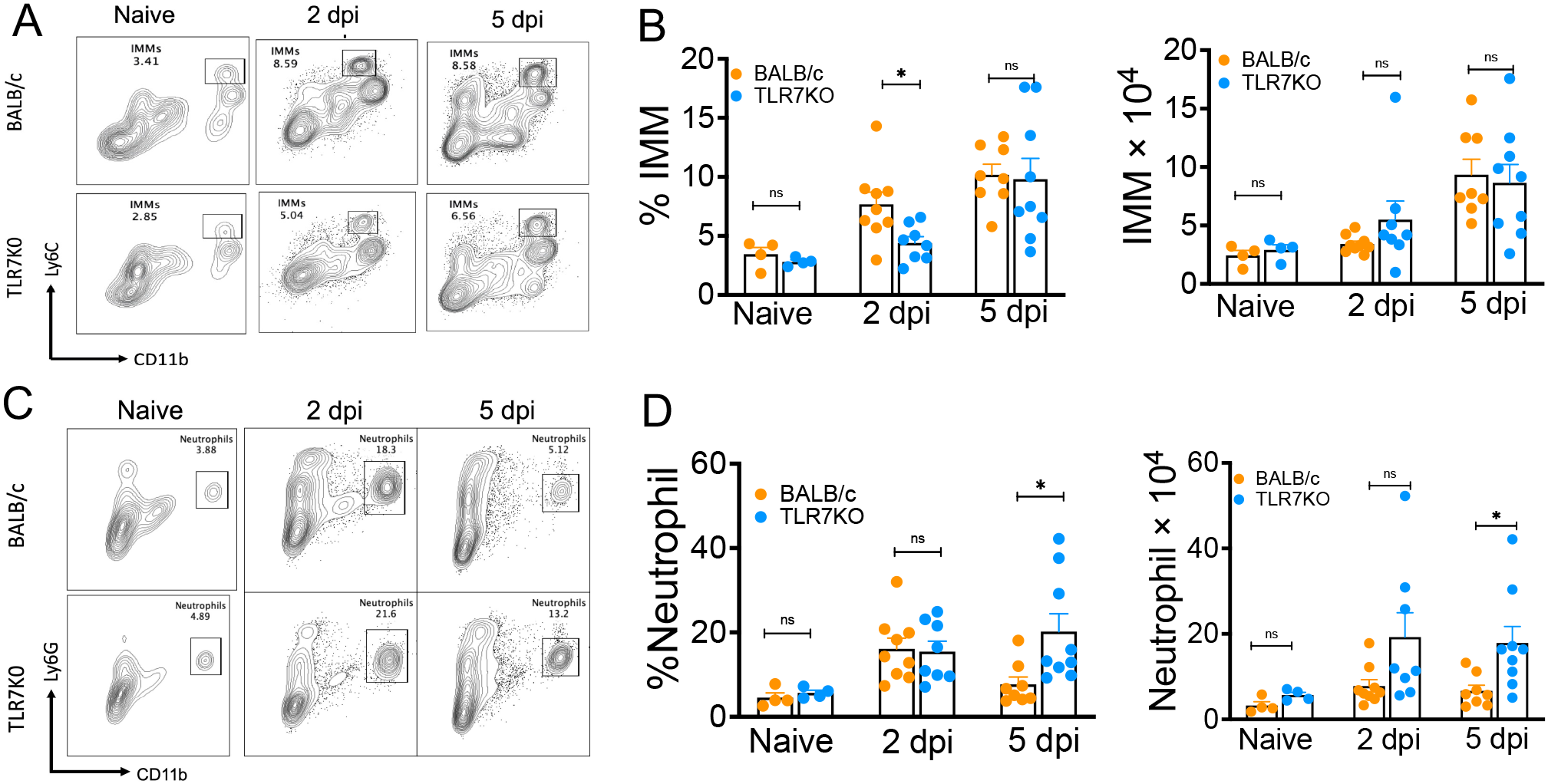
TLR7 deficiency causes differential myeloid cell accumulation in SARS-CoV-2 infected lungs. Control and TLR7KO mice were infected with 250PFU of MA-CoV-2 and lungs were harvested at indicated days post infection. Lung immune cell composition was determined by flow cytometry (A and C). FACS plots show percentage of IMM in lungs at 2 dpi and 5 dpi. (B) Bar graph shows the percentage and total IMM in the lungs at 2 dpi and 5 dpi. (C) FACS plots show percentage neutrophils in the lungs at 2 dpi and 5 dpi. (D) Bar graph shows the percentage and total neutrophils in the lungs at 2 dpi and 5 dpi. A-C) FACS data are representative two independent experiments with *n* =4-5 mice per group. B and D) results are pooled from two independent experiments with *n*=4-5 mice/group/experiment. Statistical significance was determined using Student’s *t* test with \**P* < .05 and \*\**P* < .01.

### Blocking TLR7 activity post-SARS-CoV-2 infection causes severe disease

COVID-19 studies in humans and our mouse studies show that TLR7 signaling is essential to mount an effective antiviral IFN/ISG response and provide protection during hCoV infections (Figure 1)(44, 46, 53, 54, 78-80, 91). TLR7 activation is highly pro-inflammatory, and a recent study showed pathogenic effect of TLR7 signaling during influenza virus infections with post-infection TLR7 antagonist treatment reducing inflammatory response and disease severity (92). Consequently, we evaluated whether blocking TLR7 activity after the induction of early IFN/ISG response (i.e., after day 2 post-infection) would protect mice from TLR7-induced inflammation and disease severity. We used a previously described TLR7 antagonist TSI-15-16 in our studies (93). First, we examined the ability of TLR7 antagonist to suppress TLR7 signaling. For these studies, RAW264.7 macrophages pre-treated with 20µM concentration of TLR7-antagonist were stimulation of cells with R837 (TLR7 agonist, 1µg/ml) for 6-hours in the presence of Golgi-plug. RAW264.7 cells were examined for their ability to produce TNF using flow cytometry. Our results showed a significant reduction in intracellular TNF levels compared to R837-only treated cells (Figure 6A), suggesting effective inhibition of TLR7 signaling in TSI-15-16 treated cells. Next, we treated MA-CoV-2 infected control mice with vehicle or TLR7 antagonist (dissolved in DMSO and reconstituted in PBS, 5 mg/kg body weight, via intraperitoneal route) starting 2dpi until 6 days-post infection. Control and TLR7 antagonist-treated MA-CoV-2 infected mice were monitored for morbidity and mortality for 8 days. Our results show enhanced morbidity as measured by increased loss of percent initial body weight and mortality in TLR7 antagonist-treated mice compared to control vehicle-treated groups (Figure 6B-C). These results indicate that delayed TLR7 signaling is also protective during SARS-CoV-2 infection and that blocking of TLR7 activity post-infection increases morbidity.

**Figure 6:**
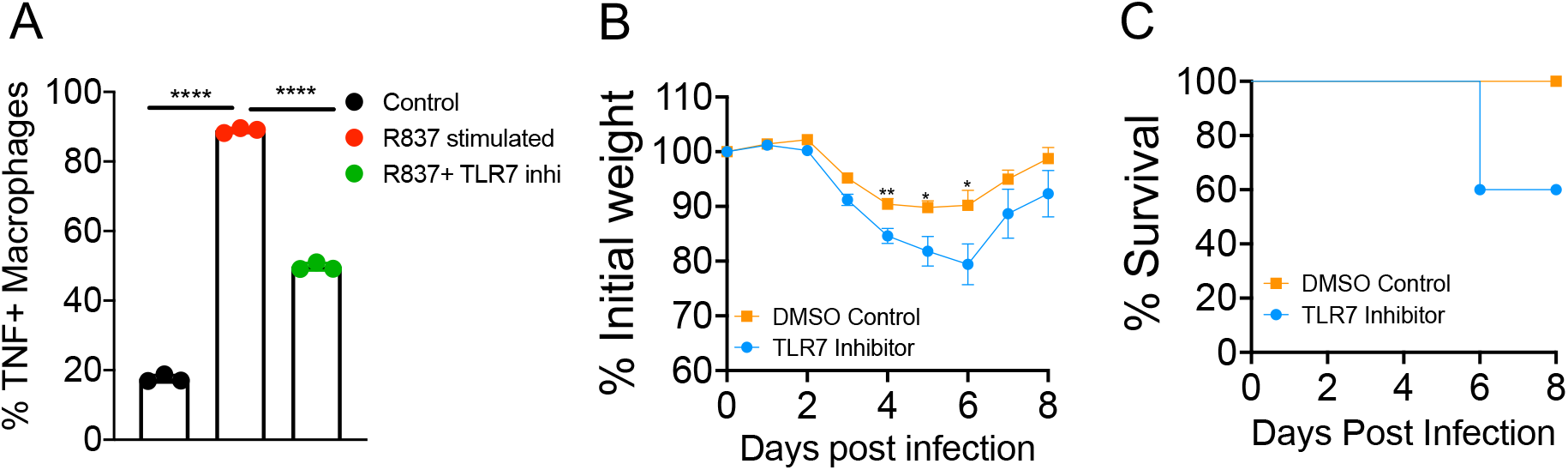
TLR7 inhibition post-infection exacerbates SARS-CoV-2-induced disease severity. A) RAW264.7 macrophages were stimulated with the TLR7 agonist R837 (1 µg/ml) with or without the TLR7 inhibitor TSI-15-16 (20 µM) for 5-6 h in the presence of Golgi-plug. Bar graph show percent TNF positive cells in control and TLR7 inhibitor-treated macrophages. B) 7–10-week-old wild type BALB/c mice were infected intranasally with 250PFU of MA-CoV-2. One group of infected mice was treated with TLR7 antagonist (5mg/kg body weight, i.p.) starting from 2dpi until 6 days post-infection while other group was treated with vehicle control. Percent initial weight (B) and survival (C) of mice were monitored for 8 days following infection. Statistical significance was determined using Student’s *t* test with \*\*\*\**P* < .001 for figure 6A, whereas One-way ANOVA with \**P* < .05 and \*\**P* < .01 was used for figure B.

## DISCUSSION

CoVs possess single-stranded RNA genome that activates TLR7/8 to induce antiviral IFN/ISG and robust inflammatory cytokine/chemokine responses (44, 46, 49, 50, 53, 54, 78-80, 83, 91). However, whether TLR7/8-indued IFN/ISG response is sufficient for host protection or TLR7/8-induced inflammatory cytokine/chemokine response promotes disease severity is not well defined. Here, we provide conclusive evidence to show that TLR7 signaling is essential for robust IFN/ISG response and is protective during SARS-CoV-2 infection in mice (Figures 1-2). These results corroborate SARS-CoV-2 studies in humans that showed severe COVID-19 in individuals with variant TLR7 or TLR7 mutation/deficiency (44, 46, 53, 54, 78-80, 91). Given similar disease outcomes in control and TLR7-deficient mice and humans, our results further demonstrate the utility of the mouse model and mouse-adapted SARS-CoV-2 to study COVID-19 pathogenesis.

Despite a central role for TLR7 signaling in the induction of robust IFN/ISG response in SARS-CoV-2 infected lungs, we did not evaluate the key cellular source of IFN-Is in MA-CoV-2 lungs. Animal studies and in-vitro data derived from murine and human PBMCs challenged/treated with SARS-CoV-2, hCOV RNA, and viral RNA mimics show that pDCs, alveolar macrophages (AMs), DCs, and monocyte-derived DCs are key sources of IFN-Is during coronavirus infection (44, 46, 53, 54, 78-80, 91). These cells are not productively infected by hCoVs and therefore it is likely that these cells produce IFNs response via abortive infection or sensing CoV ssRNA derived from dead or dying productively infected cells such as airway epithelial cells (35-39, 94). Conversely, monocyte-macrophages and neutrophils are a key source of inflammatory cytokines and chemokines (26, 84-89). The role of lung epithelial cells in SARS-COV-2-induced antiviral and inflammatory response is not known. Interestingly, lung epithelial cells express TLR7, and blocking epithelial cell TLR7 signaling using a specific inhibitor did not affect SARS-CoV-2-induced IFN-I or inflammatory cytokine response (95), suggesting a role for non-TLR7 signaling such as sensing of dsRNA by TLR3 or MDA5 in epithelial cell IFN-I response. Conversely, we showed reduced lung IFN-I response in TLR7 deficient mice intranasally transduced with adenovirus 5 encoding human dipeptidyl peptidase 4 (Ad5-hDPP4) later infected with MERS-CoV (44). Although Ad5-hDPP4 primarily transduces airway/alveolar epithelial cells, transduction and subsequent infection of AMs with MERS-CoV and AM-mediated IFN-I response activity in MERS-CoV-infected lungs cannot be ruled out. It is also likely that pDC- and likely AM-derived IFN-Is induce ISGs in epithelial cells and protect epithelial cells from SARS-CoV-2 infection (96).

Although type I and type III IFNs are protective during a variety of virus infections, their protective efficacy is dictated by the timing and magnitude of their response. These conclusions are supported by recent studies from our and other groups that showed an early IFN-I/III response relative to peak virus replication is protective during CoV and influenza virus infection, while a delayed IFN-I/III response causes severe disease (28, 44, 97, 98). Our studies further showed CoV-specific effects of IFN response in disease outcomes, as demonstrated by the detrimental role of IFNs during SARS-CoV infection, and in contrast, protective effect during mouse-CoV and MERS-CoV infection (28, 44, 46). Similar to MERS-CoV infection, we show simultaneous peak of IFNs response and virus titers during SARS-CoV-2 infection (Figure 2). Therefore, an early IFN response is likely protective during SARS-CoV-2 infection. These results are in agreement with COVID-19 studies that show severe disease in humans with genetic deficiency in IFN-I receptor signaling or autoantibodies against IFNs (59, 60, 62, 64, 65, 67, 69-71, 73). Similarly, the lack of IFNAR signaling or blocking IFNAR signaling using anti-IFNAR antibodies enhances disease severity during SARS-CoV-2 infection in animal models (84, 99-101). Our results also show a reduced IFN/ISG response in TLR7-deficient lungs (Figure 2). Therefore, it is reasonable to speculate that increased susceptibility in TLR7 deficient mice is caused by reduced IFN/ISG responses. Increased virus titers and viral sub-genomic RNA levels in TLR7 deficient lungs (Figure 4) further demonstrate the potential role of TLR7-induced IFN/ISG response in controlling SARS-CoV-2 replication in the lungs.

In addition to promoting IFN/ISG response during CoV infections, TLR7 signaling facilitates robust inflammatory cytokine and chemokine activity following viral lung infections (92, 102), including SARS-CoV-2 infection (Figure. 3). These TLR7-mediated antiviral and inflammatory responses are cell-type specific, with TLR7 inducing robust IFN response by pDCs and excessive inflammatory cytokine/ chemokine production by myeloid cells. Such inflammatory response is primarily induced following recognizing sCoV ssRNA abundant in GU- or UU-rich nucleotide sequences (49, 52, 83). Severe COVID-19 in human and animal models is associated with exuberant inflammatory cytokine and chemokine response (19, 21, 23, 26, 27, 86, 88, 103-111). Despite TLR7-dependent robust inflammatory response, it appears that TLR7-mediated IFN/ISG response is essential for early control of SARS-CoV-2 replication and host protection. Although our mouse model and human COVID-19 studies showed a protective role of TLR7 signaling during SARS-CoV-2 infection, using TLR7 inhibitors, a recent study showed a protective role of blocking TLR7 signaling during influenza virus infections (92). In contrast, and similar to TLR7 deficiency/genetic variants in humans, our results showed that blocking TLR7 signaling post-infection promotes increased morbidity and mortality, highlighting virus-specific differences in TLR7 signaling in disease outcomes following respiratory virus infections. Therefore, the use of TLR7 agonists or antagonists for therapeutic purposes requires virus-specific understanding of the role of TLR7 signaling in virus-induced antiviral and inflammatory responses.

In summary, our results show that similar to COVID-19 in humans, TLR7 signaling is protective during SARS-CoV-2 infection in mice and TLR7 induced IFN/ISG-mediated suppression of virus replication is likely critical to protect the host from lethal disease. Given similar disease outcome in TLR7 deficient humans and mice, MA-CoV-2 infected mice serve as a reliable model for studying COVID-19.

## Author Contribution

RC and RG conceptualized the study; RG, RA, RS, DC, TP, CW carried out experiments and data analysis; SM carried out histopathology evaluation; LL and TG provided reagents; RC and RG wrote the manuscript; RC, RG, RS RA, DC, TP, CW,LL, TG, and SM reviewed and edited the manuscript. RC provided mentoring and supervised experiments, data analysis, and presentation.

## Acknowledgment

The author(s) disclosed receipt of the following financial support for the research, authorship, and/or publication of this article: This work is supported in part by an institutional research fund to RC from Oklahoma State University, College of Veterinary Medicine, Oklahoma Center for Respiratory and Infectious Disease (OCRID) Centers of Biomedical Research Excellence (COBRE) grant NIH P20GM103648 and NIH AG060222. We also acknowledge OCRID/CoBRE Animal Model Core and Immunopathology Core for assistance with animal, flow cytometry, and histopathology studies.

## REFERENCES

1. http://www.who.int/csr/disease/coronavirus_infections/MERS_CoV_RA_20140613.pdf WUoM-CTfAtHaIRfA-RGLaoMAf.

2. https://covid19.who.int/ WCDC-DWw.

3. Public Health England and ISARIC, Treatment of MERS-CoV: Information for clinicians. Clinical decision-making support for treatment of MERS-CoV, https://www.gov.uk/government/uploads/system/uploads/attachment_data/file/360424/MERS_COV_information_for_clinicians_17_July.pdf.

4. WHO. 2003. Cumulative Number of Reported Probable Cases of SARS.

5. Zhu N, Zhang D, Wang W, Li X, Yang B, Song J, Zhao X, Huang B, Shi W, Lu R, Niu P, Zhan F, Ma X, Wang D, Xu W, Wu G, Gao GF, Tan W, China Novel Coronavirus I, Research T. 2020. A Novel Coronavirus from Patients with Pneumonia in China, 2019. N Engl J Med 382:727–733.

6. Peiris JS, Lai ST, Poon LL, Guan Y, Yam LY, Lim W, Nicholls J, Yee WK, Yan WW, Cheung MT, Cheng VC, Chan KH, Tsang DN, Yung RW, Ng TK, Yuen KY, group Ss. 2003. Coronavirus as a possible cause of severe acute respiratory syndrome. Lancet 361:1319–25.

7. Chen N, Zhou M, Dong X, Qu J, Gong F, Han Y, Qiu Y, Wang J, Liu Y, Wei Y, Xia J, Yu T, Zhang X, Zhang L. 2020. Epidemiological and clinical characteristics of 99 cases of 2019 novel coronavirus pneumonia in Wuhan, China: a descriptive study. Lancet 395:507–513.

8. Zhou P, Yang XL, Wang XG, Hu B, Zhang L, Zhang W, Si HR, Zhu Y, Li B, Huang CL, Chen HD, Chen J, Luo Y, Guo H, Jiang RD, Liu MQ, Chen Y, Shen XR, Wang X, Zheng XS, Zhao K, Chen QJ, Deng F, Liu LL, Yan B, Zhan FX, Wang YY, Xiao GF, Shi ZL. 2020. A pneumonia outbreak associated with a new coronavirus of probable bat origin. Nature 579:270–273.

9. Scientific Advisory Council MoH, Saudi Arabia. Infection prevention/control and managament guidelines for patients with Middle East Respiratory Syndrome Coronavirus (MERS-CoV) infection 2nd edn. http://www.moh.gov.sa/en/CCC/StaffRegulations/Corona/Documents/GuidelinesforCoronaPatients.pdf.

10. Nicholls JM, Poon LL, Lee KC, Ng WF, Lai ST, Leung CY, Chu CM, Hui PK, Mak KL, Lim W, Yan KW, Chan KH, Tsang NC, Guan Y, Yuen KY, Peiris JS. 2003. Lung pathology of fatal severe acute respiratory syndrome. Lancet 361:1773–8.

11. Arabi YM, Arifi AA, Balkhy HH, Najm H, Aldawood AS, Ghabashi A, Hawa H, Alothman A, Khaldi A, Al Raiy B. 2014. Clinical course and outcomes of critically ill patients with Middle East respiratory syndrome coronavirus infection. Ann Intern Med 160:389–97.

12. Yang Y, Guo F, Zhao W, Gu Q, Huang M, Cao Q, Shi Y, Li J, Chen J, Yan J, Jin Z, Wang X, Deng Y, Sun L, Cai H, Huang J, Zheng Y, Li W, Liu A, Chen B, Zhou M, Qiu H, Slutsky AS. 2015. Novel avian-origin influenza A (H7N9) in critically ill patients in China*. Crit Care Med 43:339–45.

13. Chu CM, Poon LL, Cheng VC, Chan KS, Hung IF, Wong MM, Chan KH, Leung WS, Tang BS, Chan VL, Ng WL, Sim TC, Ng PW, Law KI, Tse DM, Peiris JS, Yuen KY. 2004. Initial viral load and the outcomes of SARS. CMAJ 171:1349–52.

14. Hung IF, Cheng VC, Wu AK, Tang BS, Chan KH, Chu CM, Wong MM, Hui WT, Poon LL, Tse DM, Chan KS, Woo PC, Lau SK, Peiris JS, Yuen KY. 2004. Viral loads in clinical specimens and SARS manifestations. Emerg Infect Dis 10:1550–7.

15. Jacot D, Greub G, Jaton K, Opota O. 2020. Viral load of SARS-CoV-2 across patients and compared to other respiratory viruses. Microbes Infect doi:10.1016/j.micinf.2020.08.004.

16. Liu Y, Yan LM, Wan L, Xiang TX, L. A, Liu JM, Peiris M, Poon LLM, Zhang W. 2020. Viral dynamics in mild and severe cases of COVID-19. Lancet Infect Dis 20:656–657.

17. Oh MD, Park WB, Choe PG, Choi SJ, Kim JI, Chae J, Park SS, Kim EC, Oh HS, Kim EJ, Nam EY, Na SH, Kim DK, Lee SM, Song KH, Bang JH, Kim ES, Kim HB, Park SW, Kim NJ. 2016. Viral Load Kinetics of MERS Coronavirus Infection. N Engl J Med 375:1303–5.

18. Zheng S, Fan J, Yu F, Feng B, Lou B, Zou Q, Xie G, Lin S, Wang R, Yang X, Chen W, Wang Q, Zhang D, Liu Y, Gong R, Ma Z, Lu S, Xiao Y, Gu Y, Zhang J, Yao H, Xu K, Lu X, Wei G, Zhou J, Fang Q, Cai H, Qiu Y, Sheng J, Chen Y, Liang T. 2020. Viral load dynamics and disease severity in patients infected with SARS-CoV-2 in Zhejiang province, China, January-March 2020: retrospective cohort study. BMJ 369:m1443.

19. Min CK, Cheon S, Ha NY, Sohn KM, Kim Y, Aigerim A, Shin HM, Choi JY, Inn KS, Kim JH, Moon JY, Choi MS, Cho NH, Kim YS. 2016. Comparative and kinetic analysis of viral shedding and immunological responses in MERS patients representing a broad spectrum of disease severity. Sci Rep 6:25359.

20. Westblade LF, Brar G, Pinheiro LC, Paidoussis D, Rajan M, Martin P, Goyal P, Sepulveda JL, Zhang L, George G, Liu D, Whittier S, Plate M, Small CB, Rand JH, Cushing MM, Walsh TJ, Cooke J, Safford MM, Loda M, Satlin MJ. 2020. SARS-CoV-2 Viral Load Predicts Mortality in Patients with and without Cancer Who Are Hospitalized with COVID-19. Cancer Cell 38:661–671 e2.

21. van den Brand JM, Haagmans BL, van Riel D, Osterhaus AD, Kuiken T. 2014. The pathology and pathogenesis of experimental severe acute respiratory syndrome and influenza in animal models. J Comp Pathol 151:83–112.

22. Teijaro JR. 2017. Cytokine storms in infectious diseases. Semin Immunopathol 39:501–503.

23. Channappanavar R, Perlman S. 2017. Pathogenic human coronavirus infections: causes and consequences of cytokine storm and immunopathology. Semin Immunopathol 39:529–539.

24. Lau SK, Lau CC, Chan KH, Li CP, Chen H, Jin DY, Chan JF, Woo PC, Yuen KY. 2013. Delayed induction of proinflammatory cytokines and suppression of innate antiviral response by the novel Middle East respiratory syndrome coronavirus: implications for pathogenesis and treatment. J Gen Virol 94:2679–90.

25. Wang Z, Zhang A, Wan Y, Liu X, Qiu C, Xi X, Ren Y, Wang J, Dong Y, Bao M, Li L, Zhou M, Yuan S, Sun J, Zhu Z, Chen L, Li Q, Zhang Z, Zhang X, Lu S, Doherty PC, Kedzierska K, Xu J. 2014. Early hypercytokinemia is associated with interferon-induced transmembrane protein-3 dysfunction and predictive of fatal H7N9 infection. Proc Natl Acad Sci U S A 111:769–74.

26. Kuri-Cervantes L, Pampena MB, Meng W, Rosenfeld AM, Ittner CAG, Weisman AR, Agyekum R, Mathew D, Baxter AE, Vella L, Kuthuru O, Apostolidis S, Bershaw L, Dougherty J, Greenplate AR, Pattekar A, Kim J, Han N, Gouma S, Weirick ME, Arevalo CP, Bolton MJ, Goodwin EC, Anderson EM, Hensley SE, Jones TK, Mangalmurti NS, Luning Prak ET, Wherry EJ, Meyer NJ, Betts MR. 2020. Immunologic perturbations in severe COVID-19/SARS-CoV-2 infection. bioRxiv doi:10.1101/2020.05.18.101717.

27. Blanco-Melo D, Nilsson-Payant BE, Liu WC, Uhl S, Hoagland D, Moller R, Jordan TX, Oishi K, Panis M, Sachs D, Wang TT, Schwartz RE, Lim JK, Albrecht RA, tenOever BR. 2020. Imbalanced Host Response to SARS-CoV-2 Drives Development of COVID-19. Cell 181:1036–1045 e9.

28. Channappanavar R, Fehr AR, Vijay R, Mack M, Zhao J, Meyerholz DK, Perlman S. 2016. Dysregulated Type I Interferon and Inflammatory Monocyte-Macrophage Responses Cause Lethal Pneumonia in SARS-CoV-Infected Mice. Cell Host Microbe 19:181–93.

29. Arunachalam PS, Wimmers F, Mok CKP, Perera R, Scott M, Hagan T, Sigal N, Feng Y, Bristow L, Tak-Yin Tsang O, Wagh D, Coller J, Pellegrini KL, Kazmin D, Alaaeddine G, Leung WS, Chan JMC, Chik TSH, Choi CYC, Huerta C, Paine McCullough M, Lv H, Anderson E, Edupuganti S, Upadhyay AA, Bosinger SE, Maecker HT, Khatri P, Rouphael N, Peiris M, Pulendran B. 2020. Systems biological assessment of immunity to mild versus severe COVID-19 infection in humans. Science doi:10.1126/science.abc6261.

30. Hadjadj J, Yatim N, Barnabei L, Corneau A, Boussier J, Smith N, Pere H, Charbit B, Bondet V, Chenevier-Gobeaux C, Breillat P, Carlier N, Gauzit R, Morbieu C, Pene F, Marin N, Roche N, Szwebel TA, Merkling SH, Treluyer JM, Veyer D, Mouthon L, Blanc C, Tharaux PL, Rozenberg F, Fischer A, Duffy D, Rieux-Laucat F, Kerneis S, Terrier B. 2020. Impaired type I interferon activity and inflammatory responses in severe COVID-19 patients. Science 369:718–724.

31. Frieman M, Heise M, Baric R. 2008. SARS coronavirus and innate immunity. Virus Res 133:101–12.

32. Fehr AR, Perlman S. 2015. Coronaviruses: an overview of their replication and pathogenesis. Methods Mol Biol 1282:1–23.

33. Frieman M, Baric R. 2008. Mechanisms of severe acute respiratory syndrome pathogenesis and innate immunomodulation. Microbiol Mol Biol Rev 72:672–85, Table of Contents.

34. Xia H, Shi PY. 2020. Antagonism of Type I Interferon by Severe Acute Respiratory Syndrome Coronavirus 2. J Interferon Cytokine Res 40:543–548.

35. Sefik E, Qu R, Junqueira C, Kaffe E, Mirza H, Zhao J, Brewer JR, Han A, Steach HR, Israelow B, Blackburn HN, Velazquez SE, Chen YG, Halene S, Iwasaki A, Meffre E, Nussenzweig M, Lieberman J, Wilen CB, Kluger Y, Flavell RA. 2022. Inflammasome activation in infected macrophages drives COVID-19 pathology. Nature 606:585–593.

36. Junqueira C, Crespo A, Ranjbar S, de Lacerda LB, Lewandrowski M, Ingber J, Parry B, Ravid S, Clark S, Schrimpf MR, Ho F, Beakes C, Margolin J, Russell N, Kays K, Boucau J, Das Adhikari U, Vora SM, Leger V, Gehrke L, Henderson LA, Janssen E, Kwon D, Sander C, Abraham J, Goldberg MB, Wu H, Mehta G, Bell S, Goldfeld AE, Filbin MR, Lieberman J. 2022. FcgammaR-mediated SARS-CoV-2 infection of monocytes activates inflammation. Nature 606:576–584.

37. Zheng J, Wang Y, Li K, Meyerholz DK, Allamargot C, Perlman S. 2021. Severe Acute Respiratory Syndrome Coronavirus 2-Induced Immune Activation and Death of Monocyte-Derived Human Macrophages and Dendritic Cells. J Infect Dis 223:785–795.

38. Jalloh S, Olejnik J, Berrigan J, Nisa A, Suder EL, Akiyama H, Lei M, Ramaswamy S, Tyagi S, Bushkin Y, Muhlberger E, Gummuluru S. 2022. CD169-mediated restrictive SARS-CoV-2 infection of macrophages induces pro-inflammatory responses. PLoS Pathog 18:e1010479.

39. Honzke K, Obermayer B, Mache C, Fathykova D, Kessler M, Dokel S, Wyler E, Baumgardt M, Lowa A, Hoffmann K, Graff P, Schulze J, Mieth M, Hellwig K, Demir Z, Biere B, Brunotte L, Mecate-Zambrano A, Bushe J, Dohmen M, Hinze C, Elezkurtaj S, Tonnies M, Bauer TT, Eggeling S, Tran HL, Schneider P, Neudecker J, Ruckert JC, Schmidt-Ott KM, Busch J, Klauschen F, Horst D, Radbruch H, Radke J, Heppner F, Corman VM, Niemeyer D, Muller MA, Goffinet C, Mothes R, Pascual-Reguant A, Hauser AE, Beule D, Landthaler M, Ludwig S, Suttorp N, Witzenrath M, Gruber AD, Drosten C, et al. 2022. Human lungs show limited permissiveness for SARS-CoV-2 due to scarce ACE2 levels but virus-induced expansion of inflammatory macrophages. Eur Respir J 60.

40. Aboudounya MM, Heads RJ. 2021. COVID-19 and Toll-Like Receptor 4 (TLR4): SARS-CoV-2 May Bind and Activate TLR4 to Increase ACE2 Expression, Facilitating Entry and Causing Hyperinflammation. Mediators Inflamm 2021:8874339.

41. Shirato K, Kizaki T. 2021. SARS-CoV-2 spike protein S1 subunit induces pro-inflammatory responses via toll-like receptor 4 signaling in murine and human macrophages. Heliyon 7:e06187.

42. Zhao Y, Kuang M, Li J, Zhu L, Jia Z, Guo X, Hu Y, Kong J, Yin H, Wang X, You F. 2021. SARS-CoV-2 spike protein interacts with and activates TLR41. Cell Res 31:818–820.

43. Zheng M, Karki R, Williams EP, Yang D, Fitzpatrick E, Vogel P, Jonsson CB, Kanneganti TD. 2021. TLR2 senses the SARS-CoV-2 envelope protein to produce inflammatory cytokines. Nat Immunol 22:829–838.

44. Channappanavar R, Fehr AR, Zheng J, Wohlford-Lenane C, Abrahante JE, Mack M, Sompallae R, McCray PB, Jr., Meyerholz DK, Perlman S. 2019. IFN-I response timing relative to virus replication determines MERS coronavirus infection outcomes. J Clin Invest 130:3625–3639.

45. Roth-Cross JK, Bender SJ, Weiss SR. 2008. Murine coronavirus mouse hepatitis virus is recognized by MDA5 and induces type I interferon in brain macrophages/microglia. J Virol 82:9829–38.

46. Cervantes-Barragan L, Kalinke U, Zust R, Konig M, Reizis B, Lopez-Macias C, Thiel V, Ludewig B. 2009. Type I IFN-mediated protection of macrophages and dendritic cells secures control of murine coronavirus infection. J Immunol 182:1099–106.

47. Zalinger ZB, Elliott R, Rose KM, Weiss SR. 2015. MDA5 Is Critical to Host Defense during Infection with Murine Coronavirus. J Virol 89:12330–40.

48. Moreno-Eutimio MA, Lopez-Macias C, Pastelin-Palacios R. 2020. Bioinformatic analysis and identification of single-stranded RNA sequences recognized by TLR7/8 in the SARS-CoV-2, SARS-CoV, and MERS-CoV genomes. Microbes Infect 22:226–229.

49. Campbell GR, To RK, Hanna J, Spector SA. 2021. SARS-CoV-2, SARS-CoV-1, and HIV-1 derived ssRNA sequences activate the NLRP3 inflammasome in human macrophages through a non-classical pathway. iScience 24:102295.

50. Salvi V, Nguyen HO, Sozio F, Schioppa T, Gaudenzi C, Laffranchi M, Scapini P, Passari M, Barbazza I, Tiberio L, Tamassia N, Garlanda C, Del Prete A, Cassatella MA, Mantovani A, Sozzani S, Bosisio D. 2021. SARS-CoV-2-associated ssRNAs activate inflammation and immunity via TLR7/8. JCI Insight 6.

51. Forsbach A, Nemorin JG, Montino C, Muller C, Samulowitz U, Vicari AP, Jurk M, Mutwiri GK, Krieg AM, Lipford GB, Vollmer J. 2008. Identification of RNA sequence motifs stimulating sequence-specific TLR8-dependent immune responses. J Immunol 180:3729–38.

52. Heil F, Hemmi H, Hochrein H, Ampenberger F, Kirschning C, Akira S, Lipford G, Wagner H, Bauer S. 2004. Species-specific recognition of single-stranded RNA via toll-like receptor 7 and 8. Science 303:1526–9.

53. Asano T, Boisson B, Onodi F, Matuozzo D, Moncada-Velez M, Maglorius Renkilaraj MRL, Zhang P, Meertens L, Bolze A, Materna M, Korniotis S, Gervais A, Talouarn E, Bigio B, Seeleuthner Y, Bilguvar K, Zhang Y, Neehus AL, Ogishi M, Pelham SJ, Le Voyer T, Rosain J, Philippot Q, Soler-Palacin P, Colobran R, Martin-Nalda A, Riviere JG, Tandjaoui-Lambiotte Y, Chaibi K, Shahrooei M, Darazam IA, Olyaei NA, Mansouri D, Hatipoglu N, Palabiyik F, Ozcelik T, Novelli G, Novelli A, Casari G, Aiuti A, Carrera P, Bondesan S, Barzaghi F, Rovere-Querini P, Tresoldi C, Franco JL, Rojas J, Reyes LF, Bustos IG, Arias AA, et al. 2021. X-linked recessive TLR7 deficiency in ∼1% of men under 60 years old with life-threatening COVID-19. Sci Immunol 6.

54. van der Sluis RM, Cham LB, Gris-Oliver A, Gammelgaard KR, Pedersen JG, Idorn M, Ahmadov U, Hernandez SS, Cemalovic E, Godsk SH, Thyrsted J, Gunst JD, Nielsen SD, Jorgensen JJ, Bjerg TW, Laustsen A, Reinert LS, Olagnier D, Bak RO, Kjolby M, Holm CK, Tolstrup M, Paludan SR, Kristensen LS, Sogaard OS, Jakobsen MR. 2022. TLR2 and TLR7 mediate distinct immunopathological and antiviral plasmacytoid dendritic cell responses to SARS-CoV-2 infection. EMBO J 41:e109622.

55. Fallerini C, Daga S, Mantovani S, Benetti E, Picchiotti N, Francisci D, Paciosi F, Schiaroli E, Baldassarri M, Fava F, Palmieri M, Ludovisi S, Castelli F, Quiros-Roldan E, Vaghi M, Rusconi S, Siano M, Bandini M, Spiga O, Capitani K, Furini S, Mari F, Study G-CM, Renieri A, Mondelli MU, Frullanti E. 2021. Association of Toll-like receptor 7 variants with life-threatening COVID-19 disease in males: findings from a nested case-control study. Elife 10.

56. Duncan CJA, Randall RE, Hambleton S. 2021. Genetic Lesions of Type I Interferon Signalling in Human Antiviral Immunity. Trends Genet 37:46–58.

57. Matuozzo D, Talouarn E, Marchal A, Manry J, Seeleuthner Y, Zhang Y, Bolze A, Chaldebas M, Milisavljevic B, Zhang P, Gervais A, Bastard P, Asano T, Bizien L, Barzaghi F, Abolhassani H, Tayoun AA, Aiuti A, Darazam IA, Allende LM, Alonso-Arias R, Arias AA, Aytekin G, Bergman P, Bondesan S, Bryceson YT, Bustos IG, Cabrera-Marante O, Carcel S, Carrera P, Casari G, Chaibi K, Colobran R, Condino-Neto A, Covill LE, El Zein L, Flores C, Gregersen PK, Gut M, Haerynck F, Halwani R, Hancerli S, Hammarstrom L, Hatipoglu N, Karbuz A, Keles S, Kyheng C, Leon-Lopez R, Franco JL, Mansouri D, et al. 2022. Rare predicted loss-of-function variants of type I IFN immunity genes are associated with life-threatening COVID-19. medRxiv doi:10.1101/2022.10.22.22281221.

58. Zhang Q, Bastard P, Liu Z, Le Pen J, Moncada-Velez M, Chen J, Ogishi M, Sabli IKD, Hodeib S, Korol C, Rosain J, Bilguvar K, Ye J, Bolze A, Bigio B, Yang R, Arias AA, Zhou Q, Zhang Y, Onodi F, Korniotis S, Karpf L, Philippot Q, Chbihi M, Bonnet-Madin L, Dorgham K, Smith N, Schneider WM, Razooky BS, Hoffmann HH, Michailidis E, Moens L, Han JE, Lorenzo L, Bizien L, Meade P, Neehus AL, Ugurbil AC, Corneau A, Kerner G, Zhang P, Rapaport F, Seeleuthner Y, Manry J, Masson C, Schmitt Y, Schluter A, Le Voyer T, Khan T, Li J, et al. 2020. Inborn errors of type I IFN immunity in patients with life-threatening COVID-19. Science 370.

59. Abers MS, Rosen LB, Delmonte OM, Shaw E, Bastard P, Imberti L, Quaresima V, Biondi A, Bonfanti P, Castagnoli R, Casanova JL, Su HC, Notarangelo LD, Holland SM, Lionakis MS. 2021. Neutralizing type-I interferon autoantibodies are associated with delayed viral clearance and intensive care unit admission in patients with COVID-19. Immunol Cell Biol 99:917–921.

60. Bastard P, Gervais A, Le Voyer T, Rosain J, Philippot Q, Manry J, Michailidis E, Hoffmann HH, Eto S, Garcia-Prat M, Bizien L, Parra-Martinez A, Yang R, Haljasmagi L, Migaud M, Sarekannu K, Maslovskaja J, de Prost N, Tandjaoui-Lambiotte Y, Luyt CE, Amador-Borrero B, Gaudet A, Poissy J, Morel P, Richard P, Cognasse F, Troya J, Trouillet-Assant S, Belot A, Saker K, Garcon P, Riviere JG, Lagier JC, Gentile S, Rosen LB, Shaw E, Morio T, Tanaka J, Dalmau D, Tharaux PL, Sene D, Stepanian A, Megarbane B, Triantafyllia V, Fekkar A, Heath JR, Franco JL, Anaya JM, Sole-Violan J, Imberti L, et al. 2021. Autoantibodies neutralizing type I IFNs are present in ∼4% of uninfected individuals over 70 years old and account for ∼20% of COVID-19 deaths. Sci Immunol 6.

61. Bastard P, Levy R, Henriquez S, Bodemer C, Szwebel TA, Casanova JL. 2021. Interferon-beta Therapy in a Patient with Incontinentia Pigmenti and Autoantibodies against Type I IFNs Infected with SARS-CoV-2. J Clin Immunol 41:931–933.

62. Bastard P, Orlova E, Sozaeva L, Levy R, James A, Schmitt MM, Ochoa S, Kareva M, Rodina Y, Gervais A, Le Voyer T, Rosain J, Philippot Q, Neehus AL, Shaw E, Migaud M, Bizien L, Ekwall O, Berg S, Beccuti G, Ghizzoni L, Thiriez G, Pavot A, Goujard C, Fremond ML, Carter E, Rothenbuhler A, Linglart A, Mignot B, Comte A, Cheikh N, Hermine O, Breivik L, Husebye ES, Humbert S, Rohrlich P, Coaquette A, Vuoto F, Faure K, Mahlaoui N, Kotnik P, Battelino T, Trebusak Podkrajsek K, Kisand K, Ferre EMN, DiMaggio T, Rosen LB, Burbelo PD, McIntyre M, Kann NY, et al. 2021. Preexisting autoantibodies to type I IFNs underlie critical COVID-19 pneumonia in patients with APS-1. J Exp Med 218.

63. Campbell TM, Liu Z, Zhang Q, Moncada-Velez M, Covill LE, Zhang P, Alavi Darazam I, Bastard P, Bizien L, Bucciol G, Lind Enoksson S, Jouanguy E, Karabela SN, Khan T, Kendir-Demirkol Y, Arias AA, Mansouri D, Marits P, Marr N, Migeotte I, Moens L, Ozcelik T, Pellier I, Sendel A, Senoglu S, Shahrooei M, Smith CIE, Vandernoot I, Willekens K, Kart Yasar K, Effort CHG, Bergman P, Abel L, Cobat A, Casanova JL, Meyts I, Bryceson YT. 2022. Respiratory viral infections in otherwise healthy humans with inherited IRF7 deficiency. J Exp Med 219.

64. Chauvineau-Grenier A, Bastard P, Servajean A, Gervais A, Rosain J, Jouanguy E, Cobat A, Casanova JL, Rossi B. 2021. Autoantibodies neutralizing type I interferons in 20% of COVID-19 deaths in a French hospital. Res Sq doi:10.21203/rs.3.rs-915062/v1.

65. Damoiseaux J, Dotan A, Fritzler MJ, Bogdanos DP, Meroni PL, Roggenbuck D, Goldman M, Landegren N, Bastard P, Shoenfeld Y, Conrad K. 2022. Autoantibodies and SARS-CoV2 infection: The spectrum from association to clinical implication: Report of the 15th Dresden Symposium on Autoantibodies. Autoimmun Rev 21:103012.

66. de Prost N, Bastard P, Arrestier R, Fourati S, Mahevas M, Burrel S, Dorgham K, Gorochov G, Tandjaoui-Lambiotte Y, Azzaoui I, Fernandes I, Combes A, Casanova JL, Mekontso-Dessap A, Luyt CE. 2021. Plasma Exchange to Rescue Patients with Autoantibodies Against Type I Interferons and Life-Threatening COVID-19 Pneumonia. J Clin Immunol 41:536–544.

67. Koning R, Bastard P, Casanova JL, Brouwer MC, van de Beek D, with the Amsterdam UMCC-BI. 2021. Autoantibodies against type I interferons are associated with multi-organ failure in COVID-19 patients. Intensive Care Med 47:704–706.

68. Lopez J, Mommert M, Mouton W, Pizzorno A, Brengel-Pesce K, Mezidi M, Villard M, Lina B, Richard JC, Fassier JB, Cheynet V, Padey B, Duliere V, Julien T, Paul S, Bastard P, Belot A, Bal A, Casanova JL, Rosa-Calatrava M, Morfin F, Walzer T, Trouillet-Assant S. 2021. Early nasal type I IFN immunity against SARS-CoV-2 is compromised in patients with autoantibodies against type I IFNs. J Exp Med 218.

69. Solanich X, Rigo-Bonnin R, Gumucio VD, Bastard P, Rosain J, Philippot Q, Perez-Fernandez XL, Fuset-Cabanes MP, Gordillo-Benitez MA, Suarez-Cuartin G, Boza-Hernandez E, Riera-Mestre A, Parra-Martinez A, Colobran R, Antoli A, Navarro S, Rocamora-Blanch G, Framil M, Calatayud L, Corbella X, Casanova JL, Morandeira F, Sabater-Riera J. 2021. Pre-existing Autoantibodies Neutralizing High Concentrations of Type I Interferons in Almost 10% of COVID-19 Patients Admitted to Intensive Care in Barcelona. J Clin Immunol 41:1733–1744.

70. Troya J, Bastard P, Planas-Serra L, Ryan P, Ruiz M, de Carranza M, Torres J, Martinez A, Abel L, Casanova JL, Pujol A. 2021. Neutralizing Autoantibodies to Type I IFNs in >10% of Patients with Severe COVID-19 Pneumonia Hospitalized in Madrid, Spain. J Clin Immunol 41:914–922.

71. van der Wijst MGP, Vazquez SE, Hartoularos GC, Bastard P, Grant T, Bueno R, Lee DS, Greenland JR, Sun Y, Perez R, Ogorodnikov A, Ward A, Mann SA, Lynch KL, Yun C, Havlir DV, Chamie G, Marquez C, Greenhouse B, Lionakis MS, Norris PJ, Dumont LJ, Kelly K, Zhang P, Zhang Q, Gervais A, Le Voyer T, Whatley A, Si Y, Byrne A, Combes AJ, Rao AA, Song YS, Fragiadakis GK, Kangelaris K, Calfee CS, Erle DJ, Hendrickson C, Krummel MF, Woodruff PG, Langelier CR, Casanova JL, Derisi JL, Anderson MS, Ye CJ, consortium UC. 2021. Type I interferon autoantibodies are associated with systemic immune alterations in patients with COVID-19. Sci Transl Med 13:eabh2624.

72. van der Wijst MGP, Vazquez SE, Hartoularos GC, Bastard P, Grant T, Bueno R, Lee DS, Greenland JR, Sun Y, Perez R, Ogorodnikov A, Ward A, Mann SA, Lynch KL, Yun C, Havlir DV, Chamie G, Marquez C, Greenhouse B, Lionakis MS, Norris PJ, Dumont LJ, Kelly K, Zhang P, Zhang Q, Gervais A, Voyer TL, Whatley A, Si Y, Byrne A, Combes AJ, Rao AA, Song YS, consortium UC, Fragiadakis GK, Kangelaris K, Calfee CS, Erle DJ, Hendrickson C, Krummel MF, Woodruff PG, Langelier CR, Casanova JL, Derisi JL, Anderson MS, Ye CJ. 2021. Longitudinal single-cell epitope and RNA-sequencing reveals the immunological impact of type 1 interferon autoantibodies in critical COVID-19. bioRxiv doi:10.1101/2021.03.09.434529.

73. Vazquez SE, Bastard P, Kelly K, Gervais A, Norris PJ, Dumont LJ, Casanova JL, Anderson MS, DeRisi JL. 2021. Neutralizing Autoantibodies to Type I Interferons in COVID-19 Convalescent Donor Plasma. J Clin Immunol 41:1169–1171.

74. Wong LR, Zheng J, Wilhelmsen K, Li K, Ortiz ME, Schnicker NJ, Thurman A, Pezzulo AA, Szachowicz PJ, Li P, Pan R, Klumpp K, Aswad F, Rebo J, Narumiya S, Murakami M, Zuniga S, Sola I, Enjuanes L, Meyerholz DK, Fortney K, McCray PB, Jr., Perlman S. 2022. Eicosanoid signalling blockade protects middle-aged mice from severe COVID-19. Nature 605:146–151.

75. Channappanavar R, Perlman S. 2020. Evaluation of Activation and Inflammatory Activity of Myeloid Cells During Pathogenic Human Coronavirus Infection. Methods Mol Biol 2099:195–204.

76. Abolhassani H, Vosughimotlagh A, Asano T, Landegren N, Boisson B, Delavari S, Bastard P, Aranda-Guillen M, Wang Y, Zuo F, Sardh F, Marcotte H, Du L, Zhang SY, Zhang Q, Rezaei N, Kampe O, Casanova JL, Hammarstrom L, Pan-Hammarstrom Q. 2022. X-Linked TLR7 Deficiency Underlies Critical COVID-19 Pneumonia in a Male Patient with Ataxia-Telangiectasia. J Clin Immunol 42:1–9.

77. Anastassopoulou C, Gkizarioti Z, Patrinos GP, Tsakris A. 2020. Human genetic factors associated with susceptibility to SARS-CoV-2 infection and COVID-19 disease severity. Hum Genomics 14:40.

78. Butler-Laporte G, Povysil G, Kosmicki JA, Cirulli ET, Drivas T, Furini S, Saad C, Schmidt A, Olszewski P, Korotko U, Quinodoz M, Celik E, Kundu K, Walter K, Jung J, Stockwell AD, Sloofman LG, Jordan DM, Thompson RC, Del Valle D, Simons N, Cheng E, Sebra R, Schadt EE, Kim-Schulze S, Gnjatic S, Merad M, Buxbaum JD, Beckmann ND, Charney AW, Przychodzen B, Chang T, Pottinger TD, Shang N, Brand F, Fava F, Mari F, Chwialkowska K, Niemira M, Pula S, Baillie JK, Stuckey A, Salas A, Bello X, Pardo-Seco J, Gomez-Carballa A, Rivero-Calle I, Martinon-Torres F, Ganna A, Karczewski KJ, et al. 2022. Exome-wide association study to identify rare variants influencing COVID-19 outcomes: Results from the Host Genetics Initiative. PLoS Genet 18:e1010367.

79. Mantovani S, Daga S, Fallerini C, Baldassarri M, Benetti E, Picchiotti N, Fava F, Galli A, Zibellini S, Bruttini M, Palmieri M, Croci S, Amitrano S, Alaverdian D, Capitani K, Furini S, Mari F, Meloni I, Study G-CM, Frullanti E, Mondelli MU, Renieri A. 2022. Rare variants in Toll-like receptor 7 results in functional impairment and downregulation of cytokine-mediated signaling in COVID-19 patients. Genes Immun 23:51–56.

80. Solanich X, Vargas-Parra G, van der Made CI, Simons A, Schuurs-Hoeijmakers J, Antoli A, Del Valle J, Rocamora-Blanch G, Setien F, Esteller M, van Reijmersdal SV, Riera-Mestre A, Sabater-Riera J, Capella G, van de Veerdonk FL, van der Hoven B, Corbella X, Hoischen A, Lazaro C. 2021. Genetic Screening for TLR7 Variants in Young and Previously Healthy Men With Severe COVID-19. Front Immunol 12:719115.

81. van de Veerdonk FL, Netea MG. 2021. Rare variants increase the risk of severe COVID-19. Elife 10.

82. van der Made CI, Simons A, Schuurs-Hoeijmakers J, van den Heuvel G, Mantere T, Kersten S, van Deuren RC, Steehouwer M, van Reijmersdal SV, Jaeger M, Hofste T, Astuti G, Corominas Galbany J, van der Schoot V, van der Hoeven H, Hagmolen Of Ten Have W, Klijn E, van den Meer C, Fiddelaers J, de Mast Q, Bleeker-Rovers CP, Joosten LAB, Yntema HG, Gilissen C, Nelen M, van der Meer JWM, Brunner HG, Netea MG, van de Veerdonk FL, Hoischen A. 2020. Presence of Genetic Variants Among Young Men With Severe COVID-19. JAMA 324:663–673.

83. Li Y, Chen M, Cao H, Zhu Y, Zheng J, Zhou H. 2013. Extraordinary GU-rich single-strand RNA identified from SARS coronavirus contributes an excessive innate immune response. Microbes Infect 15:88–95.

84. Israelow B, Song E, Mao T, Lu P, Meir A, Liu F, Alfajaro MM, Wei J, Dong H, Homer RJ, Ring A, Wilen CB, Iwasaki A. 2020. Mouse model of SARS-CoV-2 reveals inflammatory role of type I interferon signaling. J Exp Med 217.

85. Winkler ES, Bailey AL, Kafai NM, Nair S, McCune BT, Yu J, Fox JM, Chen RE, Earnest JT, Keeler SP, Ritter JH, Kang LI, Dort S, Robichaud A, Head R, Holtzman MJ, Diamond MS. 2020. SARS-CoV-2 infection in the lungs of human ACE2 transgenic mice causes severe inflammation, immune cell infiltration, and compromised respiratory function. bioRxiv doi:10.1101/2020.07.09.196188.

86. Schulte-Schrepping J, Reusch N, Paclik D, Bassler K, Schlickeiser S, Zhang B, Kramer B, Krammer T, Brumhard S, Bonaguro L, De Domenico E, Wendisch D, Grasshoff M, Kapellos TS, Beckstette M, Pecht T, Saglam A, Dietrich O, Mei HE, Schulz AR, Conrad C, Kunkel D, Vafadarnejad E, Xu CJ, Horne A, Herbert M, Drews A, Thibeault C, Pfeiffer M, Hippenstiel S, Hocke A, Muller-Redetzky H, Heim KM, Machleidt F, Uhrig A, Bosquillon de Jarcy L, Jurgens L, Stegemann M, Glosenkamp CR, Volk HD, Goffinet C, Landthaler M, Wyler E, Georg P, Schneider M, Dang-Heine C, Neuwinger N, Kappert K, Tauber R, Corman V, et al. 2020. Severe COVID-19 Is Marked by a Dysregulated Myeloid Cell Compartment. Cell 182:1419–1440 e23.

87. Mathew D, Giles JR, Baxter AE, Oldridge DA, Greenplate AR, Wu JE, Alanio C, Kuri-Cervantes L, Pampena MB, D’Andrea K, Manne S, Chen Z, Huang YJ, Reilly JP, Weisman AR, Ittner CAG, Kuthuru O, Dougherty J, Nzingha K, Han N, Kim J, Pattekar A, Goodwin EC, Anderson EM, Weirick ME, Gouma S, Arevalo CP, Bolton MJ, Chen F, Lacey SF, Ramage H, Cherry S, Hensley SE, Apostolidis SA, Huang AC, Vella LA, Unit UPCP, Betts MR, Meyer NJ, Wherry EJ. 2020. Deep immune profiling of COVID-19 patients reveals distinct immunotypes with therapeutic implications. Science doi:10.1126/science.abc8511.

88. Silvin A, Chapuis N, Dunsmore G, Goubet AG, Dubuisson A, Derosa L, Almire C, Henon C, Kosmider O, Droin N, Rameau P, Catelain C, Alfaro A, Dussiau C, Friedrich C, Sourdeau E, Marin N, Szwebel TA, Cantin D, Mouthon L, Borderie D, Deloger M, Bredel D, Mouraud S, Drubay D, Andrieu M, Lhonneur AS, Saada V, Stoclin A, Willekens C, Pommeret F, Griscelli F, Ng LG, Zhang Z, Bost P, Amit I, Barlesi F, Marabelle A, Pene F, Gachot B, Andre F, Zitvogel L, Ginhoux F, Fontenay M, Solary E. 2020. Elevated Calprotectin and Abnormal Myeloid Cell Subsets Discriminate Severe from Mild COVID-19. Cell 182:1401–1418 e18.

89. Rosa BA, Ahmed M, Singh DK, Choreno-Parra JA, Cole J, Jimenez-Alvarez LA, Rodriguez-Reyna TS, Singh B, Gonzalez O, Carrion R, Jr., Schlesinger LS, Martin J, Zuniga J, Mitreva M, Kaushal D, Khader SA. 2021. IFN signaling and neutrophil degranulation transcriptional signatures are induced during SARS-CoV-2 infection. Commun Biol 4:290.

90. Davidson S, Crotta S, McCabe TM, Wack A. 2014. Pathogenic potential of interferon alphabeta in acute influenza infection. Nat Commun 5:3864.

91. Liu BM, Hill HR. 2020. Role of Host Immune and Inflammatory Responses in COVID-19 Cases with Underlying Primary Immunodeficiency: A Review. J Interferon Cytokine Res 40:549–554.

92. Rappe JCF, Finsterbusch K, Crotta S, Mack M, Priestnall SL, Wack A. 2021. A TLR7 antagonist restricts interferon-dependent and -independent immunopathology in a mouse model of severe influenza. J Exp Med 218.

93. Ippagunta SK, Pollock JA, Sharma N, Lin W, Chen T, Tawaratsumida K, High AA, Min J, Chen Y, Guy RK, Redecke V, Katzenellenbogen JA, Hacker H. 2018. Identification of Toll-like receptor signaling inhibitors based on selective activation of hierarchically acting signaling proteins. Sci Signal 11.

94. Thorne LG, Reuschl AK, Zuliani-Alvarez L, Whelan MVX, Turner J, Noursadeghi M, Jolly C, Towers GJ. 2021. SARS-CoV-2 sensing by RIG-I and MDA5 links epithelial infection to macrophage inflammation. EMBO J 40:e107826.

95. Bortolotti D, Gentili V, Rizzo S, Schiuma G, Beltrami S, Strazzabosco G, Fernandez M, Caccuri F, Caruso A, Rizzo R. 2021. TLR3 and TLR7 RNA Sensor Activation during SARS-CoV-2 Infection. Microorganisms 9.

96. Cervantes-Barragan L, Vanderheiden A, Royer CJ, Davis-Gardner ME, Ralfs P, Chirkova T, Anderson LJ, Grakoui A, Suthar MS. 2021. Plasmacytoid dendritic cells produce type I interferon and reduce viral replication in airway epithelial cells after SARS-CoV-2 infection. bioRxiv doi:10.1101/2021.05.12.443948.

97. Dijkman R, Verma AK, Selvaraj M, Ghimire R, Gad HH, Hartmann R, More S, Perlman S, Thiel V, Channappanavar R. 2022. Effective Interferon Lambda Treatment Regimen To Control Lethal MERS-CoV Infection in Mice. J Virol 96:e0036422.

98. Major J, Crotta S, Llorian M, McCabe TM, Gad HH, Priestnall SL, Hartmann R, Wack A. 2020. Type I and III interferons disrupt lung epithelial repair during recovery from viral infection. Science 369:712–717.

99. Chong Z, Karl CE, Halfmann PJ, Kawaoka Y, Winkler ES, Keeler SP, Holtzman MJ, Yu J, Diamond MS. 2022. Nasally delivered interferon-lambda protects mice against infection by SARS-CoV-2 variants including Omicron. Cell Rep 39:110799.

100. Ogger PP, Garcia Martin M, Michalaki C, Zhou J, Brown JC, D. Y, Miah KM, Habib O, Hyde SC, Gill DR, Barclay WS, Johansson C. 2022. Type I interferon receptor signalling deficiency results in dysregulated innate immune responses to SARS-CoV-2 in mice. Eur J Immunol 52:1768–1775.

101. Uddin MB, Liang Y, Shao S, Palani S, McKelvey M, Weaver SC, Sun K. 2022. Type I IFN Signaling Protects Mice from Lethal SARS-CoV-2 Neuroinvasion. Immunohorizons 6:716–721.

102. Goritzka M, Makris S, Kausar F, Durant LR, Pereira C, Kumagai Y, Culley FJ, Mack M, Akira S, Johansson C. 2015. Alveolar macrophage-derived type I interferons orchestrate innate immunity to RSV through recruitment of antiviral monocytes. J Exp Med 212:699–714.

103. Cameron MJ, Ran L, Xu L, Danesh A, Bermejo-Martin JF, Cameron CM, Muller MP, Gold WL, Richardson SE, Poutanen SM, Willey BM, DeVries ME, Fang Y, Seneviratne C, Bosinger SE, Persad D, Wilkinson P, Greller LD, Somogyi R, Humar A, Keshavjee S, Louie M, Loeb MB, Brunton J, McGeer AJ, Canadian SRN, Kelvin DJ. 2007. Interferon-mediated immunopathological events are associated with atypical innate and adaptive immune responses in patients with severe acute respiratory syndrome. J Virol 81:8692–706.

104. Bermejo-Martin JF, Martin-Loeches I, Rello J, Anton A, Almansa R, Xu L, Lopez-Campos G, Pumarola T, Ran L, Ramirez P, Banner D, Ng DC, Socias L, Loza A, Andaluz D, Maravi E, Gomez-Sanchez MJ, Gordon M, Gallegos MC, Fernandez V, Aldunate S, Leon C, Merino P, Blanco J, Martin-Sanchez F, Rico L, Varillas D, Iglesias V, Marcos MA, Gandia F, Bobillo F, Nogueira B, Rojo S, Resino S, Castro C, Ortiz de Lejarazu R, Kelvin D. 2010. Host adaptive immunity deficiency in severe pandemic influenza. Crit Care 14:R167.

105. Teijaro JR, Walsh KB, Cahalan S, Fremgen DM, Roberts E, Scott F, Martinborough E, Peach R, Oldstone MB, Rosen H. 2011. Endothelial cells are central orchestrators of cytokine amplification during influenza virus infection. Cell 146:980–91.

106. Zheng Y, Liu X, Le W, Xie L, Li H, Wen W, Wang S, Ma S, Huang Z, Ye J, Shi W, Ye Y, Liu Z, Song M, Zhang W, Han JJ, Belmonte JCI, Xiao C, Qu J, Wang H, Liu GH, Su W. 2020. A human circulating immune cell landscape in aging and COVID-19. Protein Cell doi:10.1007/s13238-020-00762-2.

107. de Jong MD, Hien TT. 2006. Avian influenza A (H5N1). J Clin Virol 35:2–13.

108. de Jong MD, Simmons CP, Thanh TT, Hien VM, Smith GJ, Chau TN, Hoang DM, Chau NV, Khanh TH, Dong VC, Qui PT, Cam BV, Ha do Q, Guan Y, Peiris JS, Chinh NT, Hien TT, Farrar J. 2006. Fatal outcome of human influenza A (H5N1) is associated with high viral load and hypercytokinemia. Nat Med 12:1203–7.

109. Lee JS, Park S, Jeong HW, Ahn JY, Choi SJ, Lee H, Choi B, Nam SK, Sa M, Kwon JS, Jeong SJ, Lee HK, Park SH, Park SH, Choi JY, Kim SH, Jung I, Shin EC. 2020. Immunophenotyping of COVID-19 and influenza highlights the role of type I interferons in development of severe COVID-19. Sci Immunol 5.

110. Veras FP, Pontelli MC, Silva CM, Toller-Kawahisa JE, de Lima M, Nascimento DC, Schneider AH, Caetite D, Tavares LA, Paiva IM, Rosales R, Colon D, Martins R, Castro IA, Almeida GM, Lopes MIF, Benatti MN, Bonjorno LP, Giannini MC, Luppino-Assad R, Almeida SL, Vilar F, Santana R, Bollela VR, Auxiliadora-Martins M, Borges M, Miranda CH, Pazin-Filho A, da Silva LLP, Cunha LD, Zamboni DS, Dal-Pizzol F, Leiria LO, Siyuan L, Batah S, Fabro A, Mauad T, Dolhnikoff M, Duarte-Neto A, Saldiva P, Cunha TM, Alves-Filho JC, Arruda E, Louzada-Junior P, Oliveira RD, Cunha FQ. 2020. SARS-CoV-2-triggered neutrophil extracellular traps mediate COVID-19 pathology. J Exp Med 217.

111. Zuo Y, Yalavarthi S, Shi H, Gockman K, Zuo M, Madison JA, Blair C, Weber A, Barnes BJ, Egeblad M, Woods RJ, Kanthi Y, Knight JS. 2020. Neutrophil extracellular traps in COVID-19. JCI Insight 5.

